# Arktos: a simple tool for the design of polyhedral DNA nanostructures

**DOI:** 10.1101/2024.02.07.576791

**Authors:** Harshitha Balaji, Anish Hemanth Samprathi, Rakshita Sukruth Kolipakala, Pushya Pradeep, Deepesh Nagarajan

**Affiliations:** Department of Biotechnology, M.S. Ramaiah University of Applied Sciences, Bangalore - 560054, India; Department of Biotechnology, Fergusson College (Autonomous), Pune - 411004, India; Department of Microbiology, St. Xavier’s College, Mumbai - 400001, India

## Abstract

DNA nanostructures are a class of self-assembling nanomaterials with a wide range of potential applications in biomedicine and nanotechnology. The history of DNA nanotechnology can be traced back to the 1980s with the development of simple DNA polyhedra using either human intuition or simple algorithms. Today the field is dominated by DNA origami constructs to such an extent that the original algorithms used to design non-origami nanostructures have been lost. In this work we describe Arktos: an algorithm developed to design simple DNA polyhedra without the use of DNA origami. Arktos designs sequences predicted to fold into a desired structure using simulated annealing optimization. As a proof-of-concept, we used Arktos to design a simple DNA tetrahedron. The generated oligonucleotide sequences were synthesized and experimentally validated via polyacrylamide gel electrophoresis, indicating that they fold into the desired structure. These results demonstrate that Arktos can be used to design custom DNA polyhedra as per the needs of the research community.

## Introduction

DNA nanocages and nanostructures are a class of self-assembling nanomaterials that have been developed over the past several decades. These structures are formed by the hybridization of complementary DNA strands, and they can be designed to have a variety of shapes and sizes. DNA nanocages and nanostructures have a wide range of potential applications in biomedicine and nanotechnology, including drug delivery, biosensing, and tissue engineering^1^.

The history of DNA nanotechnology can be traced back to the 1980s, when Ned Seeman and his colleagues developed the first DNA nanostructures^2^. Seeman’s early work focused on the design of DNA junctions, which are the basic building blocks of DNA nanostructures. Early DNA nanotechnology involved the design of nanostructures using self-assembling DNA oligonucleotides. Early accomplishments using this approach include the design of a simple DNA cube^3^, a truncated octahedron^4^, a tetrahedron^5^, a truncated bipyramid, and a wide array of prisms^6^.

In the early 1990s, Paul Rothemund developed the DNA origami technique, which allows for the creation of complex DNA nanostructures with arbitrary shapes^1^. DNA origami uses short DNA oligonucleotides, or ‘staples’ that bind to and fold a long single stranded ‘scaffold’ into any desired shape. Currently, it is possible to create any arbitrary 2-dimensional or 3-dimensional shape using DNA origami. Examples include its use to create a 2D ‘smiley face’^1^, a polyhedral mesh resembling the Stanford bunny^7^, and nanocages possessing an assortment of sizes, shapes, and functions^8–16^.

Despite its advantages, DNA origami requires the synthesis of hundreds of staples, making it an expensive approach. Furthermore, the requirement for a scaffold and staples makes the synthesis and assembly of a DNA origami-based nanostructure very difficult *in vivo*. Simple oligonucleotide-based DNA nanostructures would be preferable under such restraints.

There exists a plethora of software used for the creation of DNA nanostructures, such as caDNAno^17^, Tiamat^18^, CanDo^19–21^, VHelix^7^, and OxDNA^22^. However, most of these tools are optimized for the design of DNA origami-based nanostructures. Furthermore, all of the tools mentioned rely on graphical user interfaces, making the high-throughput programmatic design of DNA nanostructures difficult. Early DNA oligomer-based nanostructures were either designed by hand or using tools that are no longer available.

In this work we describe Arktos: a simple tool optimized for the design of DNA oligomer-based nonstructures. Arktos reduces DNA nanostructures to simple graphical abstractions containing nodes and edges. Arktos performs both positive and negative design via simulated annealing while designing sequences to fit the inputted nanostructure topologies. We computationally validated Arktos by benchmarking it against sequences designed to fold into tetrahedra^5^. We experimentally validated Arktos by designing a simple DNA tetrahedron containing 4 strands that were seen to assemble on a PAGE gel. Furthermore, Arktos can be run on a linux terminal, making the programatic generation of a large variety of DNA nanostructures possible. We have made Arktos freely available through this work (Supplementary Protocol S1) as a github repository (https://github.com/1337deepesh/Arktos). We hope Arktos will be of use to the research community.

## Results and Discussion

We describe the Arktos algorithm, its computational validation, and its experimental validation via the creation of a simple DNA tetrahedron in the sections below.

### Algorithm design

Arktos attempts to design simple DNA polyhedra using positive and negative design. DNA polyhedra are reduced to graphical abstractions for input (Figure 2B,C). ssDNA-ssDNA interactions that form dsDNA are represented as edges, or connections.

Positive design involves trying to design on-target matches (base-pair complementarity) for every matching strand. Negative design ensures that off-target matches that arise by chance in unpaired strands are eliminated. Arktos does this using a simple scoring function:

Let:

*LM* = Length of a single off-target match (length *≥*4) between a pair of strands (including self-pairings).

*M* = Total number off-target matches (length *≥*4) between a pair of strands (including self-pairings).

*n* = Total number nucleotides in all strands.

*N* = Total number of strands.

Then:

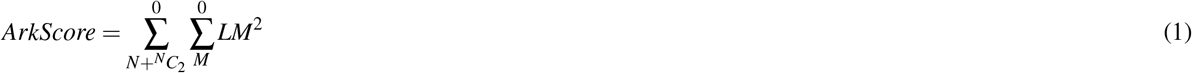

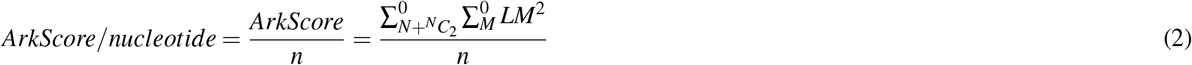

For a system of *N* strands, there are *N* +^*N*^ *C*_2_ total strand pairings (including self pairings at off-target regions). A summation of the squares of all off-target matches of sequence lengths *≥*4 across all *N* +^*N*^ *C*_2_ strands is the ArkScore (Equation 1). A sequence length of *≥*4 was chosen as the minimum threshold for a mismatch since mismatch lengths *≤*3 only transiently bind and will not affect the foldability of our designed polyhedron. A square term is added to disproportionately penalize off-target matches of greater lengths.

The ArkScore will be zero or positive, with lower scores indicating fewer off-target matches. By design, it can never be negative. A simulated annealing algorithm^23^is used to minimize the ArkScore in order to maximize folding into the designed polyhedral structure. A normalized Arkscore (ArkScore/nucleotide, Equation 2) can be used to compare DNA polyhedra with different numbers of strands and sequence lengths.

Designed sequences are outputted in the fasta format, but with ‘|’ breaks between different complementary regions (nodes) within a strand. The Arktos script, along with full details on the input and output formats, can be found in Protocol S1 and in our Github repository (https://github.com/1337deepesh/Arktos).

### Algorithm validation

We performed two exercises to computationally validate Arktos: Firstly, we evaluated the scalability of Arktos. Here ‘scalability’ denotes the ability of Arktos to output consistent scores irrespective of the size of the DNA nanostructure under design. We evaluated scalability using a simple DNA trimer with extendable strands (Figure 1A). Strands A, B, and C form interactions through ‘nodes’, or short complementary DNA stretches. DNA trimers containing 10*s*2000 nucleotide nodes were chosen for scalability evaluation (Figure 1B). We observed that the ArkScore/nucleotide remains fairly constant for designed DNA nanostructures within the same magnitude of size, increasing by only 0.01877 units per nucleotide. This increase is expected as larger DNA nanostructures inherently possesses a greater number of off-target strand permutations. When comparing DNA nanostructures across different orders of magnitude of size, a normalized score (ArkScore_*OM*_, Equation 3) can be used that compensates for this increase.

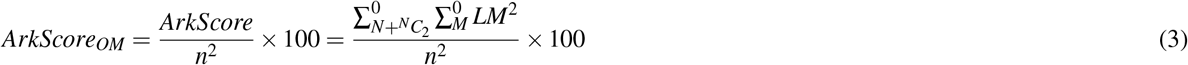

Secondly, we benchmarked Arktos against experimentally-validated DNA tetrahedra designed by Goodman *et al*.^5^. In a preliminary communication^24^, Goodman *et al*. mentioned that the sequences were designed to “minimize the strength of undesirable interactions”, but did not describe any method. We scored DNA tetrahedra designed by Goodman *et al*. using Arktos. Further, we used Arktos to redesign the sequences of Goodman *et al*. tetrahedra, while keeping their overall topology intact. We found a statistically significant difference between these undesigned and redesigned scores (Table 1, Figure 1C). Redesigned tetrahedra possessed significantly lower ArkScore/nucleotide values compared to undesigned Goodman *et al*. tetrahedra.

**Figure 1.**
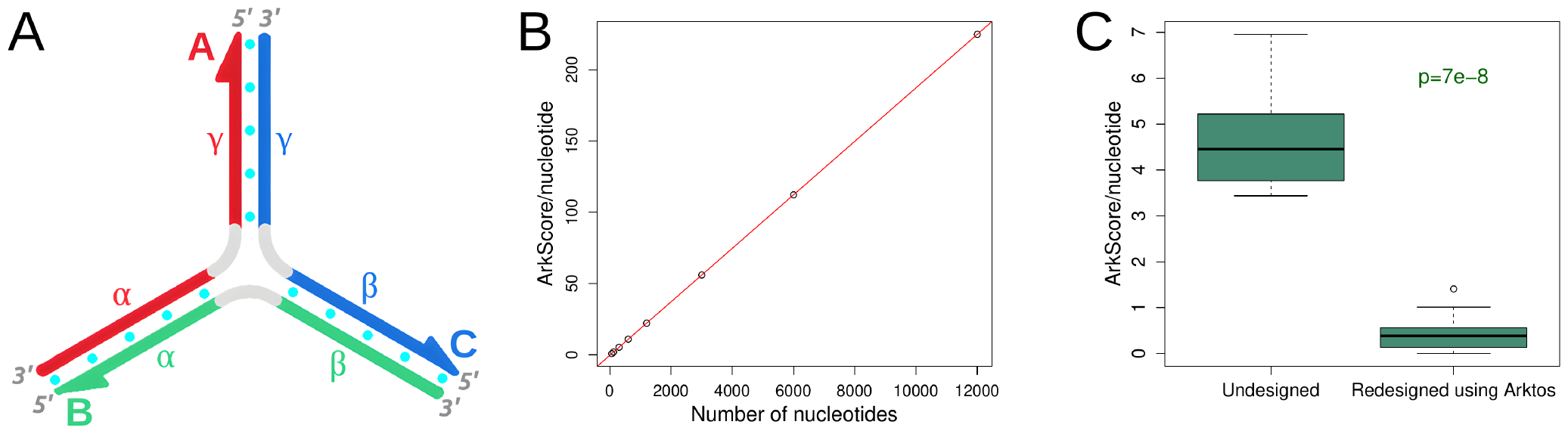
Computational evaluation of Arktos. **(A)** A DNA trimer containing strands A, B, and C was used for testing design scalability. Each strand can be subdivided into nodes. For example, strand A possesses nodes *α*, hinge, and *γ*. Hinges are single unpaired adenine residues and are colored grey. Inter-strand base pairing is represented with cyan dots. The DNA trimer depicted here possesses 5 base pairs per complementary region (10 base pairs per strand). However, complementary regions containing 10, 20, 50, 100, 200, 500, 1000, and 2000 base pairs were used for evaluation. **(B)** ArkScore/nucleotide (Equations 1 and 2) increases linearly with an increasing number of nucleotides (sum from all 3 strands). ArkScore/nucleotide increases by 0.01877 units per nucleotide. A normalized score (ArkScore/nucleotide^2^*×*100) can be used while comparing DNA nanostructures across different orders of magnitude in size. However, ArkScore/nucleotide can be used to compare DNA nanostructures of the same magnitude in size with negligible error. **(C)** Benchmarking of Arktos against previously designed DNA tetrahedra^5^. Firstly, we scored the existing tetrahedra using Arktos. Secondly, we redesigned their sequence using Arktos, while keeping their overall topology intact. The Arktos scores of redesigned tetrahedra were significantly lower (*p* = 7 *×* 10^*−*8^, Welch 2-sample T-test) compared to previously designed tetrahedra merely scored using Arktos. A detailed score breakdown is provided in Table 1.

**Figure 2.**
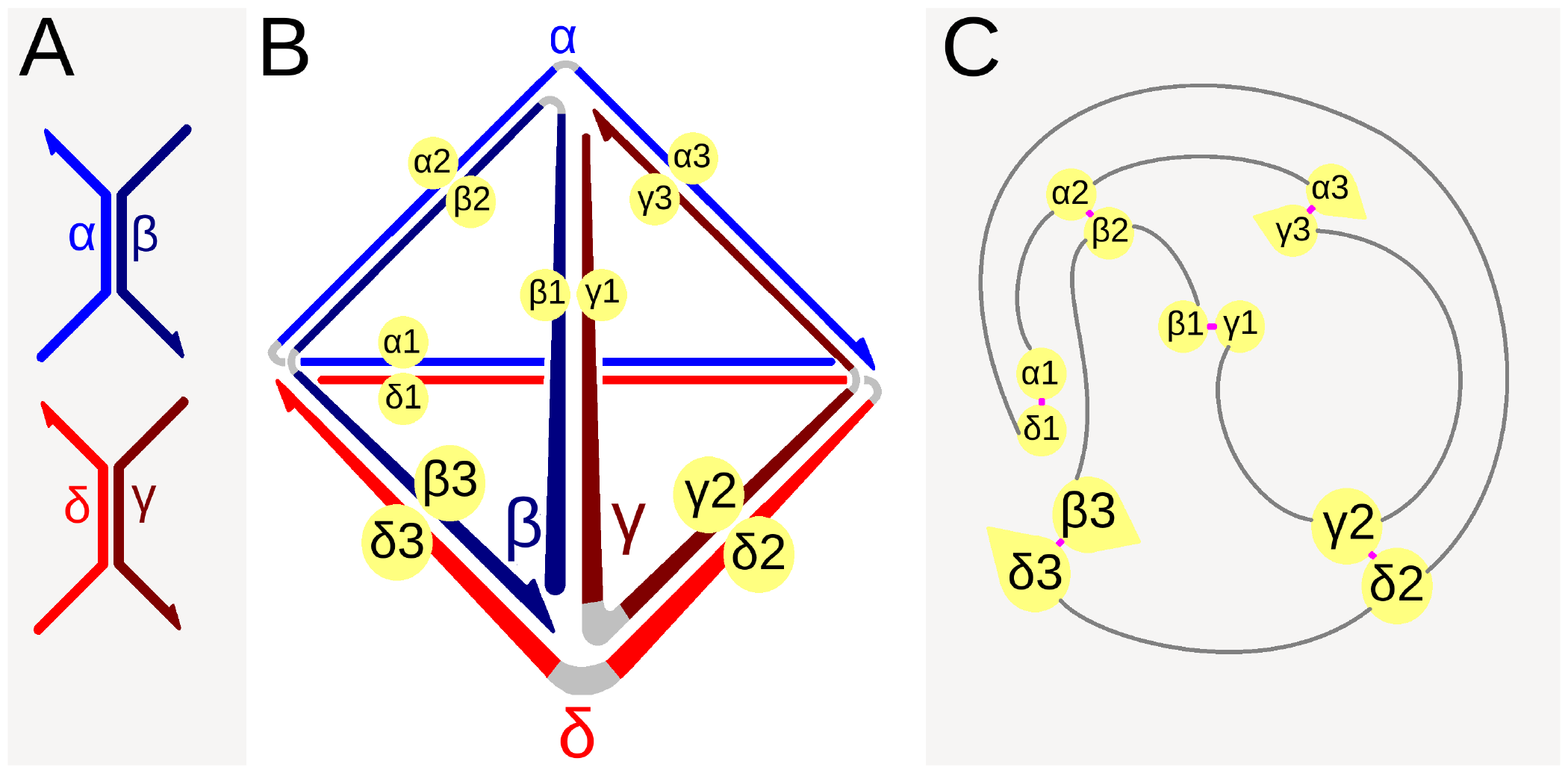
Design of a simple DNA tetrahedron to validate Arktos. 4 DNA oligomers named *α, β, γ, δ* were designed using Arktos and synthesized. Their sequences are provided in Table 2 **(A)** Strand-pairings in isolation. DNA oligomers *α −β* and *γ −δ* are shown as pairs to help the reader understand their placement in the full DNA assembly. **(B)** Strand-pairings within the DNA tetrahedron. Individual strands are divided into ‘nodes’ based on their strand-pairings. Strand *α*, for example, is divided into nodes *α*1, hinge1, *α*2, hinge2, and *α*3. Gray regions indicate hinges. All hinges are composed of lone unpaired adenine nucleotides. **(C)** Reducing the DNA tetrahedron to a graphical representation for input into Arktos. Individual nodes are now represented as points with ‘edges’ connecting to other nodes. Magenta edges represent designed complementary strand pairings. Grey edges represent hinges that are always connected to 2 nodes on either end. Any arbitrary polyhedral shape is reducible to a graphical representation.

**Table 1.**
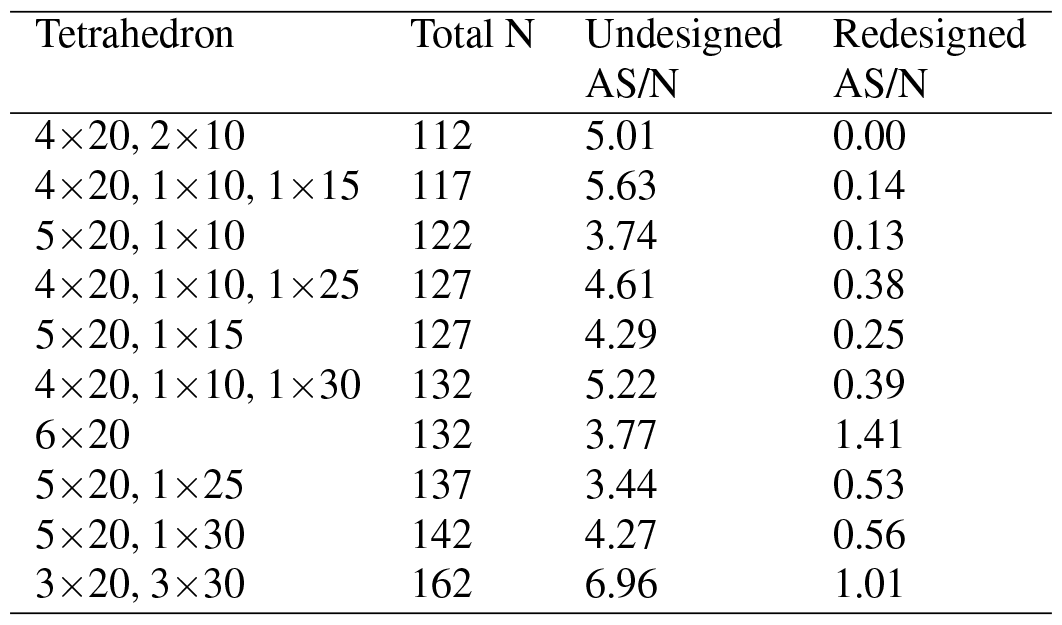
A comparison of ArkScore/nucleotide (AS/N) for previously designed DNA tetrahedra^5^ redesigned using Arktos vs. previously designed DNA tetra simply scored using Arktos (undesigned). The tetrahedral architecture is provided in the first column. 4*×*20, 2*×*10, for example means the designed tetrahedron contained 4 edges (ds DNA strand pairings) of 20 nucleotides and 2 edges of 10 nucleotides (6 edges total, see Figure 2B). ‘Total N’ refers to the total number of nucleotides in all DNA strands of a given tetrahedron. This value includes both paired edges and unpaired hinges. The ArkScore/nucleotide of Redesigned DNA tetrahedra is significantly lower than the ArkScore/nucleotide of undesigned tetrahedra.

These results indicate that while Goodman *et al*. tetrahedra were experimentally observed to rapidly fold into their desired structure, the methods used by Goodman *et al*. may not be scalable for larger DNA polyhedra. Larger polyhedra require larger DNA sequences, which will inherently possesses a greater number of off-target strand permutations. This makes the design of larger DNA polyhedra or nanostructures from DNA oligonucleotides inherently more difficult. Therefore Arktos is expected to outperform the method used by Goodman *et al*. for larger DNA nanostructures, but further experimental validation via the synthesis and characterization of such structures is required to confirm.

### Experimental validation

Using Arktos, we designed a simple DNA tetrahedron possessing an edge-length of 31 nucleotides with mono-adenyl hinges. The entire structure possessed 4 strands containing 92 nucleotides (Table 2), with the entire structure possessing 368 nucleotides in total. The topological design of this structure is provided in Figure 2. Figure 2 represents the 4 DNA strands: *α, β, γ* and *δ* and their expected sub-assemblies. For example, the central regions of strands *α* and *β* are designed to be complementary, while the 5’ and 3’ regions are designed to be complementary to the 5’ and 3’ regions of strands *γ* and *δ*. The entire assembly is provided in Figure 2B. Each strand is subdivided into ‘nodes’ designed to be complementary to nodes on other strands. This assembly is reduced to a graphical abstraction (Figure 2C) containing only nodes and edges for input into Arktos.

**Table 2.**
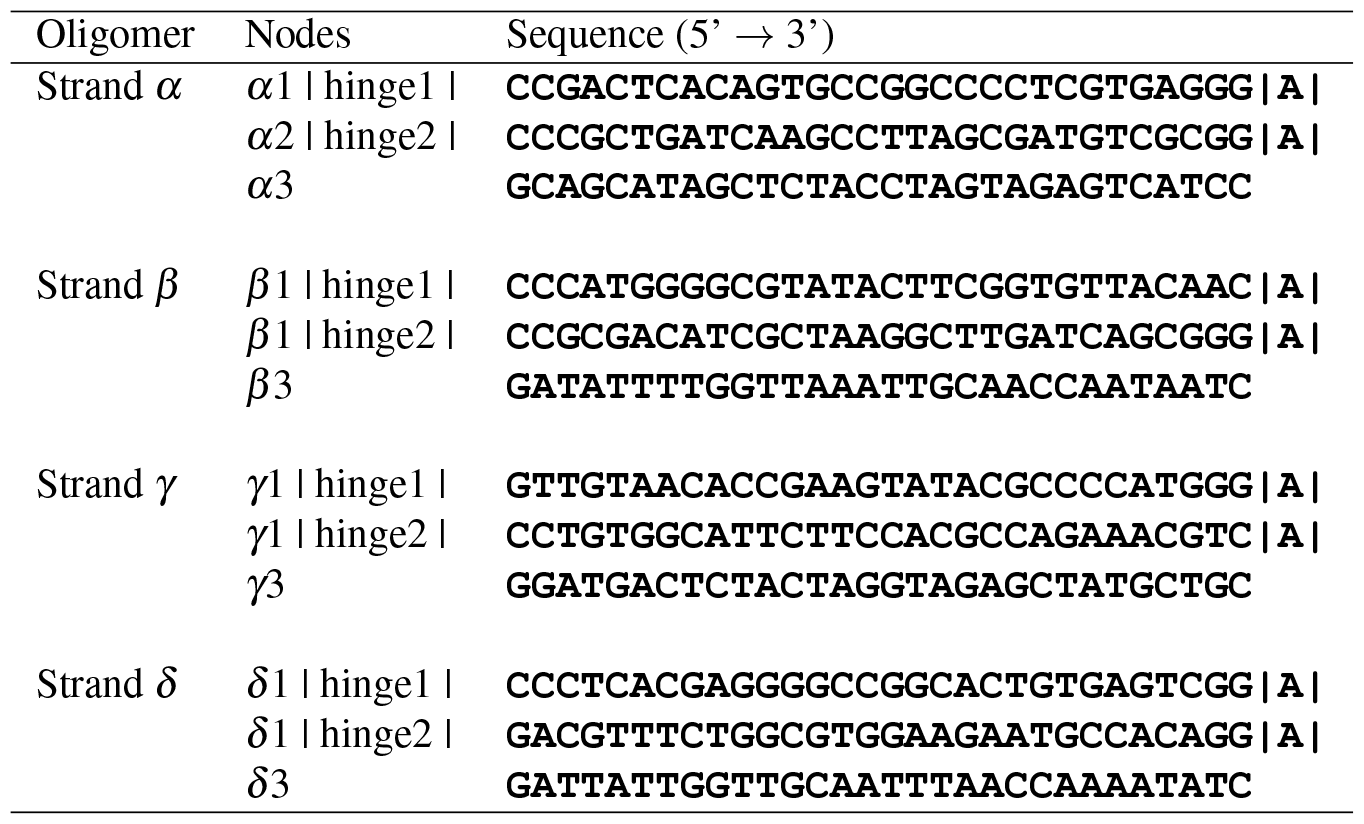
Nucleotide sequences for all 4 DNA strands designed using Arktos. These strands were designed to form a tetrahedral nanostructure upon assembly. ‘|’ breaks between sequences represent the starts/ends of nodes (including hinges).

Arktos uses simulated annealing^23^ to optimize designed sequences to minimize off-target matches. As simulated annealing is a stochastic heuristic process, it will generate different outputs in close approximation to the global minima. Arktos was therefore run 1000 times using the same input graph, and the output sequences possessing the lowest ArkScore were selected for oligomer synthesis.

The 4 designed oligomers were mixed in 1:1 or 1:1:1:1 stoichiometric ratios and annealed as described in the Methods section. The annealed products were visualized on a PAGE gel (Figure 3). We observed single bands for the annealed products of strands *α* and *β* (lane 2) as well as strands *γ* and *δ* (lane 3). This indicates that these strand-pairs dimerize, as depicted in Figure 2A. Strands *α, β, γ* and *δ* were also annealed in a one-pot approach, with the annealed product also displaying a single band (lane 5) higher along the gel than the *α −β* (lane 2) and *γ −δ* (lane 3) bands, indicating the assembly of an ordered nanostructure. Finally, strands *α, β, γ* and *δ* display aggregation when run on the gel without annealing (lane 4), indicating that annealing is required to form an ordered nanostructure.

**Figure 3.**
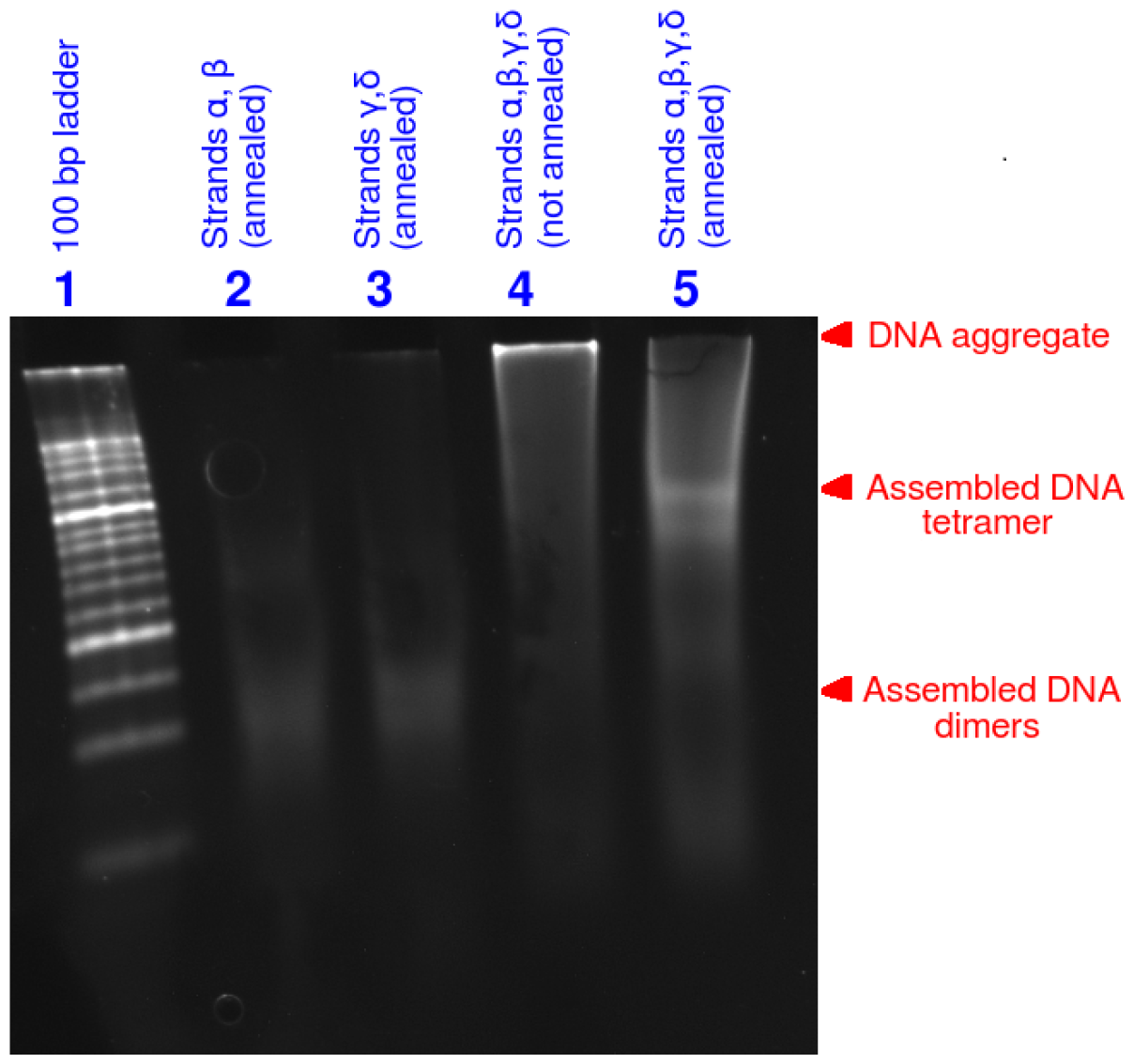
Polyacrylamide gel electrophoresis. confirms that DNA strands *α, β, γ*, and *δ* assemble into ordered structures. **Lane 1**: 100 bp ladder. **Lane 2**: Strands *α* and *β* annealed together. They form a dimer as expected. **Lane 3**: Strands *γ* and *δ* annealed together. They form a dimer as expected. **Lane 4**: Strands *α, β, γ*, and *δ* run without annealing. The strands aggregate, showing that annealing is necessary to form an ordered structure. **Lane 5**: Strands *α, β, γ*, and *δ* run after annealing. The strands display an ordered assembly expected to be tetrahedral. However, further experiments are required to confirm its structure.

These results strongly indicate that strands *α, β, γ* and *δ* folded into the desired DNA tetrahedron. However, further experimentation using cryo-electron microscopy, atomic force microscopy, or X-ray diffraction are required to confirm its shape.

In conclusion, we have developed Arktos: a simulated annealing-based algorithm to design DNA nanostructures from DNA oligomers. We have computationally validated the algorithm against existing DNA tetrahedra designed by Goodman *et al*.^5^. We have also experimentally validated Arktos by using it to design a simple DNA tetrahedron. We have made Arktos freely available hope it will find applications within the research community.

## Methods

### Oligomer synthesis

The sequences of our 4 designed DNA oligomers (Table 2) were synthesized by Biotech Desk Pvt. Ltd. (Bangalore, India).

### Oligomer annealing

All DNA oligomers were dissolved in TE buffer at a stock concentration of 170 *µ*M. Oligomers were diluted in folding buffer which contained 50 mM Tris-Cl (Tris-Base, Sigma 77861; HCl, Fisher Scientific 29507) pH 8.0, 12.5 mM MgCl2 (EMPLURA 105833) and 0.2 mM EDTA (SRL 35888) to give final concentration of 10 *µ*M which was then put through the annealing process and used in further steps^16^. Oligomers were heated to 90^*°*^C for 5 minutes and then cooled to 4^*°*^C in steps of 0.1^*°*^C every 5 seconds using a thermocycler to allow the DNA strands to anneal. The assembled structures were then stored at -20^*°*^C until ready for analysis.

### Nanostructure electrophoresis

Before loading the samples onto the gel, they were thawed at room temperature for 20 minutes. An 8% polyacrylamide gel (29:1) was prepared in 1*×* TAE/Mg^2+^ buffer and used to analyze the formation of the self-assembled nanostructures. 20 *µ*L of each sample was loaded into a well on the gel, along with 5 *µ*L of loading dye containing 0.003% bromophenol blue and 60% glycerol in 1*×* TAE/Mg^2+^ buffer. A 100 bp DNA Ladder (Invitrogen™15628019) was used as molecular weight marker in the first lane.

The gel was run at a constant voltage of 100V for 80 minutes in a cold room of 4^*°*^C. The cold temperature helps to prevent the DNA sequences from denaturing. The gel was then stained with GelRed (Biotium 41003) stain diluted to 3*×* in 0.1 M NaCl solution for 30 minutes on a gel rocker. GelRed is a fluorescent dye that binds to DNA. The gel was then visualized in using BioRad Image Labs 6.1.

## Acknowledgements

This project was funded by the M.S. Ramaiah University of Applied Sciences, Office of Research and Innovation (grant number: ORI-SG/FLAHS/001/2023. We thank Abhinav Banerjee and Dr. Mahipal Ganji (Department of Biochemistry, Indian Institute of Science, Bangalore) for their invaluable help during the experimental validation of our designed DNA tetrahedron.

## Author contributions statement

D.N. conceived and developed the algorithm and Arktos software package. P.P. and D.N. conceived the experiments, H.B., A.S. and R.K. conducted the experiments. All authors analyzed the experimental data. All authors reviewed the manuscript.

## Supplementary material

Protocol S1: The Arktos software package. The package is also available as a GitHub repository (https://github.com/1337deepesh/Arktos).

## References

1. Rothemund, P. W. Folding dna to create nanoscale shapes and patterns. Nature 440, 297–302 (2006).

2. Seeman, N. C. An overview of structural dna nanotechnology. Mol. biotechnology 37, 246–257 (2007).

3. Chen, J. & Seeman, N. C. Synthesis from dna of a molecule with the connectivity of a cube. Nature 350, 631–633 (1991).

4. Zhang, Y. & Seeman, N. C. Construction of a dna-truncated octahedron. J. Am. Chem. Soc. 116, 1661–1669 (1994).

5. Goodman, R. P. et al. Rapid chiral assembly of rigid dna building blocks for molecular nanofabrication. Science 310, 1661–1665 (2005).

6. Nie, Z. et al. Self-assembly of dna nanoprisms with only two component strands. Chem. communications 49, 2807–2809 (2013).

7. Benson, E. et al. Dna rendering of polyhedral meshes at the nanoscale. Nature 523, 441–444 (2015).

8. Zhao, Z., Jacovetty, E. L., Liu, Y. & Yan, H. Encapsulation of gold nanoparticles in a dna origami cage. Angewandte Chemie 123, 2089–2092 (2011).

9. Chandrasekaran, A. R. & Levchenko, O. Dna nanocages. Chem. Mater. 28, 5569–5581 (2016).

10. Zhang, Q. et al. Dna origami as an in vivo drug delivery vehicle for cancer therapy. ACS nano 8, 6633–6643 (2014).

11. Ijas, H., Hakaste, I., Shen, B., Kostiainen, M. A. & Linko, V. Reconfigurable dna origami nanocapsule for ph-controlled encapsulation and display of cargo. ACS nano 13, 5959–5967 (2019).

12. Luo, X., Chidchob, P., Rahbani, J. F. & Sleiman, H. F. Encapsulation of gold nanoparticles into dna minimal cages for 3d-anisotropic functionalization and assembly. Small 14, 1702660 (2018).

13. Udomprasert, A. & Kangsamaksin, T. Dna origami applications in cancer therapy. Cancer science 108, 1535–1543 (2017).

14. Han, X. et al. Multivalent aptamer-modified tetrahedral dna nanocage demonstrates high selectivity and safety for anti-tumor therapy. Nanoscale 11, 339–347 (2019).

15. Xing, S. et al. Constructing higher-order dna nanoarchitectures with highly purified dna nanocages. ACS applied materials & interfaces 7, 13174–13179 (2015).

16. Banerjee, A., Anand, M., Kalita, S. & Ganji, M. Single-molecule analysis of dna base-stacking energetics using patterned dna nanostructures. Nat. Nanotechnol. 1–9 (2023).

17. Douglas, S. M. et al. Rapid prototyping of 3d dna-origami shapes with cadnano. Nucleic acids research 37, 5001–5006 (2009).

18. Williams, S. et al. Tiamat: a three-dimensional editing tool for complex dna structures. In DNA Computing: 14th International Meeting on DNA Computing, DNA 14, Prague, Czech Republic, June 2-9, 2008. Revised Selected Papers 14, 90–101 (Springer, 2009).

19. Castro, C. E. et al. A primer to scaffolded dna origami. Nat. methods 8, 221–229 (2011).

20. Kim, D.-N., Kilchherr, F., Dietz, H. & Bathe, M. Quantitative prediction of 3d solution shape and flexibility of nucleic acid nanostructures. Nucleic acids research 40, 2862–2868 (2012).

21. Pan, K., Bricker, W. P., Ratanalert, S. & Bathe, M. Structure and conformational dynamics of scaffolded dna origami nanoparticles. Nucleic acids research 45, 6284–6298 (2017).

22. Poppleton, E., Romero, R., Mallya, A., Rovigatti, L. & Šulc, P. Oxdna. org: a public webserver for coarse-grained simulations of dna and rna nanostructures. Nucleic acids research 49, W491–W498 (2021).

23. Bertsimas, D. & Tsitsiklis, J. Simulated annealing. Stat. science 8, 10–15 (1993).

24. Goodman, R. P., Berry, R. M. & Turberfield, A. J. The single-step synthesis of a dna tetrahedron. Chem. Commun 1372–1373 (2004).

